# Ciliary IFT88 safeguards coordinated epiphyseal vascularisation, resorption and ossification from disruptive physiological mechanical forces

**DOI:** 10.1101/2021.08.06.455437

**Authors:** C. R. Coveney, H. J. Samvelyan, J. Miotla-Zarebska, J. Carnegie, E. Chang, C. J. Corrin, T. Coveney, B. Stott, I. Parisi, C. Duarte, T. L. Vincent, K. A. Staines, A.K.T. Wann

**Affiliations:** Centre for OA pathogenesis Versus Arthritis, The Kennedy Institute of Rheumatology, University of Oxford, Oxford, United Kingdom; School of Pharmacy and Biomolecular Sciences, University of Brighton, Brighton, UK

## Abstract

In the musculoskeletal system, appropriate cell and tissue responses to mechanical force delineate morphogenesis and ensure lifelong health. Despite this, how mechanical cues are integrated into biological programmes remains unclear. Primary cilia are microtubule-based organelles that tune a range of cell activities, including signalling cascades activated or modulated, by extracellular biophysical cues. Here, we demonstrate that the inducible, cartilage-specific deletion of Intraflagellar transport protein 88 (IFT88), which reduces ciliation in the adolescent mouse growth plate (GP), uncouples chondrocyte differentiation from cartilage resorption and mineralisation in a mechano-dependent manner. Targeting IFT88, inhibits hypertrophic chondrocyte VEGF expression, vascular recruitment, osteoclastic activity and the replacement of cartilage with bone. These effects are largely restricted to peripheral tibial regions beneath the load-bearing compartments of the knee. Increases in physiological loading, in control mice, also impairs ossification in the peripheral GP, mimicking the effects of IFT88 deletion. Strikingly, limb immobilisation rescues disrupted VEGF and restores epiphyseal dynamics in *Ift88*cKO mice. These data indicate, that during this pivotal phase in adolescent skeletal maturation that defines the cessation of growth, ciliary IFT88 protects the coordinated ossification of the growth plate from an otherwise disruptive heterogeneity of physiological mechanical forces.

## Introduction

All biological processes take place in the presence of mechanical forces (1). Biophysical environmental cues must be assimilated into pre-programmed genetic plans; cells and the extracellular matrix (ECM) collectively integrate mechanical forces to orchestrate tissue mechanoadaptations befitting developmental period and location. The creation, maturation and homeostasis of the musculoskeletal (MSK) system depends upon the regulated integration of, and balanced response to, mechanical and biological cues. How forces are translated into appropriate mechano-adaptations, at tissue level, remains to be understood and is a challenging question to address.

The primary cilium has been proposed to play a central role in cellular mechanotransduction (2–6). However, the mechanism by which cilia transduce or influence the cellular response to mechanical force in health and disease is still debated (7–11). A singular, microtubule-based organelle assembled by the vast majority of cell types, the cilium is a well-established nexus for the transduction of external cues, acting as a nanoscale scaffold for the regulation of multiple signalling pathways, including growth factor signalling (12–15). The ciliopathies, congenital disorders associated with mutations to ciliary-associated genes or biology, have a well-described MSK subset (16), demonstrating the fundamental importance of the primary cilium in human skeletal development. The developmental depletion of key ciliary genes in the mouse (17–23) results in impaired growth, and premature epiphyseal fusion, when the growth plate (GP), the cartilaginous template for long bone formation, fuses to become bone early. Far less is known about ciliary influence in adulthood, but ciliary IFT88 remains influential in post-natal articular cartilage (24).

Longitudinal bone growth is underpinned by endochondral ossification (EO), a carefully coordinated process of cell and tissue differentiation, that ultimately results in GP cartilage being replaced by bone. Elongation requires the GP to be organised into columns of chondrocytes, continuously supplied, throughout growth, by a stem cell niche (25, 26). Progeny of this niche undergo proliferation, enlargement through hypertrophic differentiation (27) and ultimately either apoptosis or transdifferentiation (28), all the while secreting and remodelling a regionally specialised extracellular matrix. Thus, a highly organised sequence of cellular and extracellular signalling events enables dynamic, almost simultaneous mineralisation and resorption of cartilage, vascular invasion and the creation of bone. During EO, complex gradients of growth factor signalling coordinate differentiation of cells and matrix. One example of signalling underpinning this programme of differentiation, is the Indian hedgehog (Ihh)-Parathyroid hormone-related protein (PThrP) feedback loop, which acts to balance proliferation and hypertrophic differentiation (29–33). In a similar fashion to targeting of cilia in early development, as cilia are central regulators of Hedgehog (Hh) signalling, disruption of this loop by genetic perturbation, results in accelerated GP closure (12, 13, 15). Comparatively to EO, the signalling events underlying fusion of the GP, the abrupt discontinuation of EO demarcating the cessation of growth, are poorly understood.

Both Hh signalling, PThrP signalling, and their downstream effects have themselves been previously demonstrated to be mechano-regulated. For example, hydrostatic strain applied to GP chondrocytes results in increased Ihh signalling and proliferation (34). Indeed, either by modulation of the expression of ligands, receptors or by their release from sequestration within the matrix, growth factor signalling in cartilage and bone is highly mechano-regulated (35–37). A number of studies illustrate the importance of mechanics in animal models of bone growth. In the absence of mechanical forces exerted by muscular contraction, proliferation decreased in the GP of embryonic chicks (38, 39). Tissue mechanics are also required for the intercalation of growth plate chondrocytes to affect extension (40, 41). Despite the importance of mechanotransduction to skeletal development, health and disease, the cellular and molecular components that might comprise a system that supports mechanical homeostasis in cartilage and many other tissues, analogous to the bone mechanostat originally proposed by Frost (42), remain elusive.

We hypothesised that, IFT88, and by extension the primary cilium, maintains profound influence in the post-natal growth plate. We show ciliary IFT88 plays an instrumental role in coordinating adolescent epiphyseal biology *in vivo*. We propose that cilia protect the carefully orchestrated cessation of growth, from otherwise disruptive mechanical forces and offer a new paradigm for the role of cilia in tissue mechanotransduction during morphogenesis.

## Results

### Deletion of IFT88 in the juvenile and adolescent growth plate inhibits endochondral ossification and growth plate closure

To delete *Ift88,* in a cartilage-specific, inducible, manner *AggrecanCreER^T2^;Ift88^fl/fl^* (*Ift88* cKO) mice were generated. To assess efficacy of Cre recombination in growth plate (GP) chondrocytes, the *AggrecanCreER^T2^* was crossed with a TdTomato reporter line. Effective Cre recombination was identified in many, but not all, GP chondrocyte columns following tamoxifen administration (n=3) (Figure. 1A). Chondrocyte columns expressing Tdtomato were evenly spread throughout the GP, with no bias to particular regions. By the point of analysis, 2 weeks post-tamoxifen, cells within the Primary Spongiosa also expressed Tdtomato. Immunohistochemical (IHC) staining of cryosections from 10-week-old mice, enabled visualisation of cilia *in situ* (Figure 1B, Supplementary figure 1A) and indicated a ~20% reduction in cilia prevalence in GP chondrocytes (Figure 1B, ****p<0.0001, Fisher’s exact test, contingency data shown in Supplementary Figure. 1B, n=4 in each group). Whilst in the mouse the GP never fully fuses, on approach to skeletal maturity, the rate of longitudinal bone growth decreases between 4 and 10 weeks of age, and tibial GP length reduces with age from approximately 0.26mm to 0.04mm (Figure. 1C) indicative of GP closure. Tamoxifen was administered to control and *AggrecanCreER^T2^;Ift88^fl/fl^* mice (*Ift88* cKO) at 4, or 6, or 8 weeks of age (Figure. 1D). GP lengths were analysed two weeks later, using MicroCT images of whole knee joints, taking the mean of 8 length measurements across the full width (Supplementary Figure. 1C). Analysis revealed, deletion of IFT88 resulted in statistically significantly longer GP, compared with controls (****p<0.0001, two-way ANOVA, Figure. 1E). Whilst variance increased, GP length in *AggrecanCreER^T2^;Ift88^fl/fl^*, remained similar to that of control mice at the age tamoxifen was administered (two weeks prior) across all timepoints. Thus, GP narrowing during each of these periods was effectively abolished. Though not the focus of this study it was clear that the bone in the tibial diaphysis, particularly at 6 weeks of age, appeared increased in density. Analysis of the region of bone directly beneath the GP revealed an increase in BV/TV at 6 weeks of age (Supplementary Figure. 2A). Strikingly, elongated cartilaginous GP in *AggrecanCreER^T2^;Ift88^fl/fl^* were characterised by large, regions with little or no mineral density that were largely restricted to one or both sides of the tibia. (Figure.1F).

**Figure 1.**
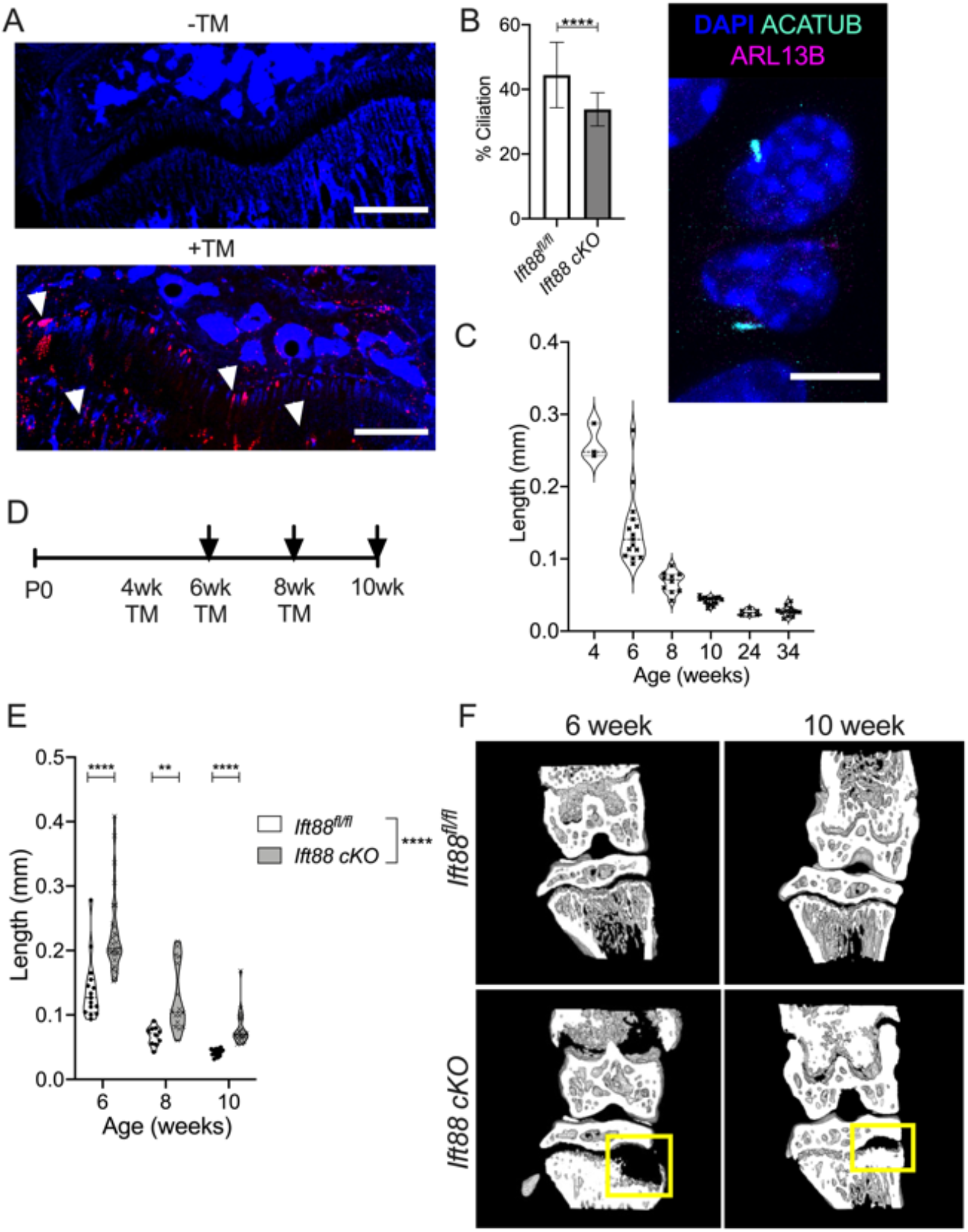
Deletion of IFT88 reduces growth plate ciliation and inhibits growth plate closure. **A,** Cryosections of knee joints counterstained with nuclear DAPI, were taken from 6-week-old AggrecanCreER^T2^;TdTomato mice that received tamoxifen (TM) at 4 weeks of age (scale bar =500um). White arrows point to chondrocyte populations exhibiting TdTomato reporter activity. **B,** Percentage ciliation in 10-week-old GP in AggrecanCreER^T2^;Ift88^fl/fl^ and Ift88^fl/fl^ control mice 2 weeks after tamoxifen. **** p=<0.0001, Fisher’s Exact test, contingency data shown in supplementary Figure 1B, Error bars shown are S.D. Image depicts primary cilia staining in GP chondrocytes in situ. **C,** Violin plots depict GP length of control animals (also treated with tamoxifen) at 4, 6, 8, 10, 24, and 34 weeks of age. **D,** Schematic indicates age of tamoxifen administration (TM) and collection (arrows). **E,** Violin plots depict GP lengths of control (Ift88^fl/fl^) and AggrecanCreER^T2^;Ift88^fl/fl^ (cKO) mice at 6, 8, and 10 weeks of age. **F,** Partial 3D construction of MicroCT scans at 6 and 10 weeks of age. Points represent mean GP length per animal in violin plots Genotype effect analysed by two-way ANOVA, pairwise analysis by unpaired t-tests corrected for multiplicity and using a 1% FDR, *p<0.05, ****p<0.0001.

### IFT88 deletion inhibits ossification of the peripheral growth plate

MicroCT images suggested the effects of IFT88 deletion were restricted to the peripheral regions of the GP, directly below the load-bearing articular surfaces of the knee (Figure. 2A), while comparatively the central region of the GP, had narrowed normally through ossification. Considering only the 8-10 week period, in order to focus on GP fusion processes and avoid the confounding effect of tibial widening at earlier timepoints (Supplementary Figure. 2B), maximum GP length measurements were taken in lateral, central and medial regions of control and *AggrecanCreER^T2^;Ift88^fl/fl^* mice and plotted relative to control animals (Figure. 2B). This analysis revealed the largest effects were observed in the medial peripheral region of the GP, where GP length was twice that of controls. Comparatively modest effects on GP length were measurable in the lateral region, whilst only very small, but nevertheless statistically significant, differences were observed centrally (Figure. 2B). Von Kossa staining also indicated disruption to mineralisation and trabecular organisation beneath the peripheral regions of failed ossification in *AggrecanCreER^T2^;Ift88^fl/fl^* mice (Figure 2C). Previous studies investigating GP closure describe bone bridging events associated with heterogenous local tissue mechanical stresses (43). Fewer and lower density bone bridges were observed in *AggrecanCreER^T2^;Ift88^fl/fl^* mice compared with controls (Figure. 2D). This reduction in bridging was again particularly striking on the medial side of the limb (Figure. 2E).

**Figure 2.**
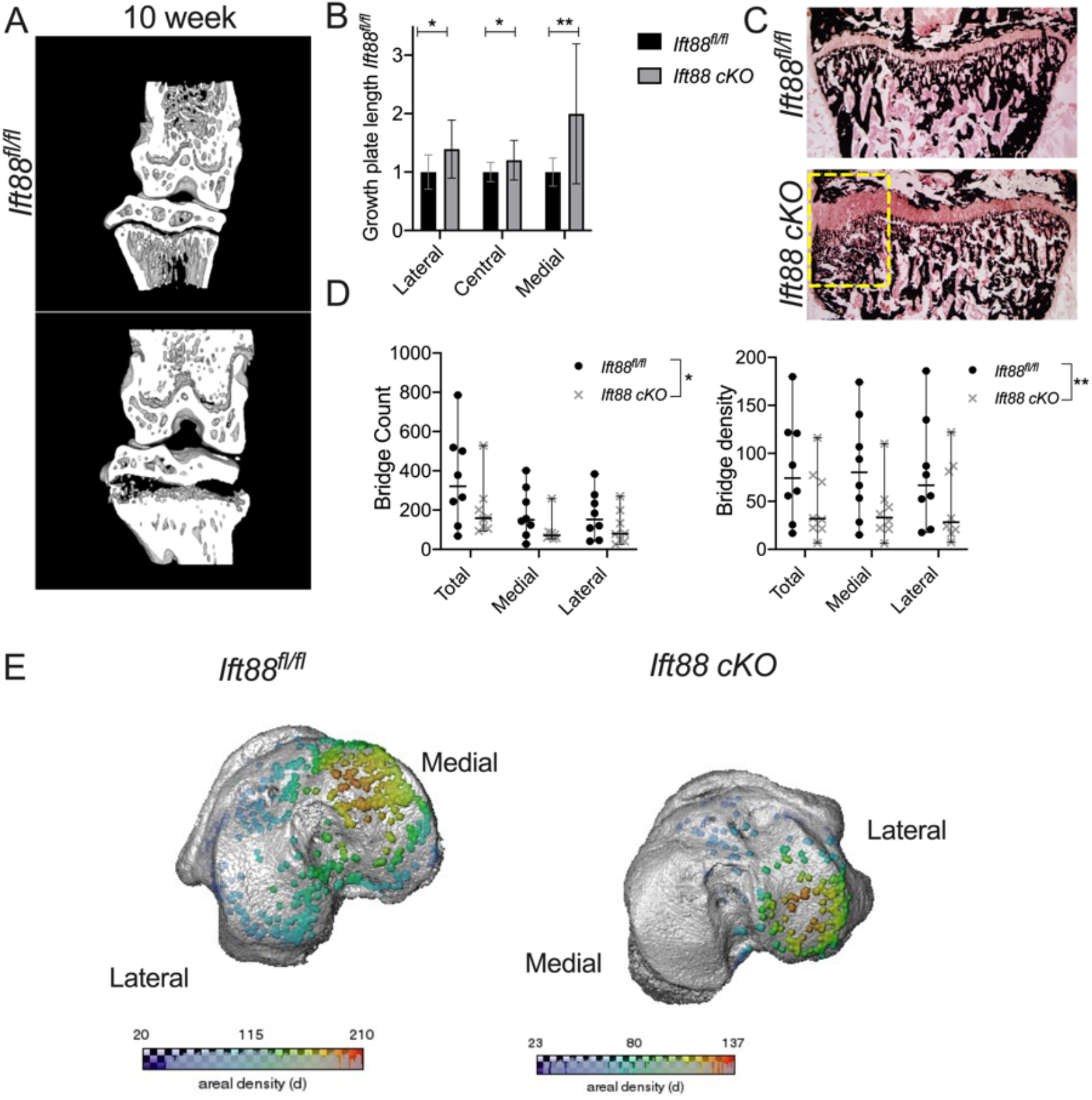
IFT88 deletion inhibits peripheral growth plate ossification. **A,** MicroCT partial 3D construction of AggrecanCreER^T2^;Ift88^fl/fl^ mice at 10 weeks of age. **B,** Maximum growth plate lengths taken from the lateral, medial and central sections of the GP of AggrecanCreER^T2^;Ift88^fl/fl^ animals, normalised to the Ift88^fl/fl^. Analysis by pairwise unpaired t-tests corrected for multiplicity, FDR 1%, *p=<0.05, **p=<0.01 Ift88^fl/fl^ n=17; AggrecanCreER^T2^;Ift88^fl/fl^ n=19. **C**, Von Kossa staining of Ift88^fl/fl^ and AggrecanCreER^T2^;Ift88^fl/fl^. Label highlights region of medial bone with disorganised trabeculae. **D,** Number of bridges and bridge density in control and AggrecanCreER^T2^;Ift88^fl/fl^. Points represent median value per animal. Analysed by two-way ANOVA, *p<0.05, **p<0.01, Ift88^fl/fl^ n=8; AggrecanCreER^T2^;Ift88^fl/fl^ n=8. **E**, 3D representation mapping GP bridges across tibial articular surfaces of the knee. Colour scale indicates the density of the bridges.

### Increased physiological loading disrupts peripheral growth plate closure

Previous modelling has indicated heterogeneity of physiological mechanical stresses across the width of the GP during limb loading (43, 44). Given the peripheral, most pronounced on the medial side, pattern to the failed ossification in *AggrecanCreER^T2^;Ift88^fl/fl^* mice, we hypothesised that coordinated epiphyseal closure is sensitive to the depletion of IFT88, due to a critical role for cilia in mechanosensation/transduction, as has been previously proposed (45, 46). Therefore, we surmised GP narrowing in control mice, between 8 and 10 weeks of age, would be sensitive to acute changes in limb loading. First, we tested the effect of removing mechanical input to the adolescent GP hypothesising this may inhibit GP dynamics in a similar manner to that observed upon depletion of cilia. Double neurectomy was performed on the right hind limb at 8 weeks of age. Cutting both the femoral and sciatic nerves rendered the right hind limb incapable of weight bearing (off-loaded), whilst the left (contralateral) became the predominant weight bearing hind limb taking increased load by means of compensation. MicroCT revealed GP in off-loaded limbs of 10 weeks old *Ift88^fl/fl^* control mice were not strikingly different when compared with naïve *Ift88^fl/fl^* control mice (Figure 3A, quantified in 3D) and relative narrowing, across the width of the limb, was uniform. However, the contralateral limbs of operated animals exhibited similar bi-lateral regions of failed ossification to that observed in naïve *AggrecanCreER^T2^;Ift88^fl/fl^* mice (Figure. 3A, right-hand image), with an associated increase in average GP length when compared with paired off-loaded limbs (Figure 3B). In order to further investigate whether increases, rather than decreases, to physiological loading, disrupt coordinated GP ossification, mice were given access to free wheel exercise between 8 and 10 weeks of age. At 10 weeks of age exercised *Ift88^fl/fl^* control mice also exhibited inhibition of GP closure in the periphery, again often most pronounced on the medial side (Figure. 3C, relative quantification of mean GP length in Figure. 3D). Von Kossa staining confirmed a failure of mineralisation and alterations to bone architecture beneath in control mice after two weeks of wheel exercise (Figure 3E). Von Kossa indicated the inhibitory effects on mineralisation were greatest on the medial side, but that the mineralised architecture beneath the lateral plateaus was also altered.

**Figure 3.**
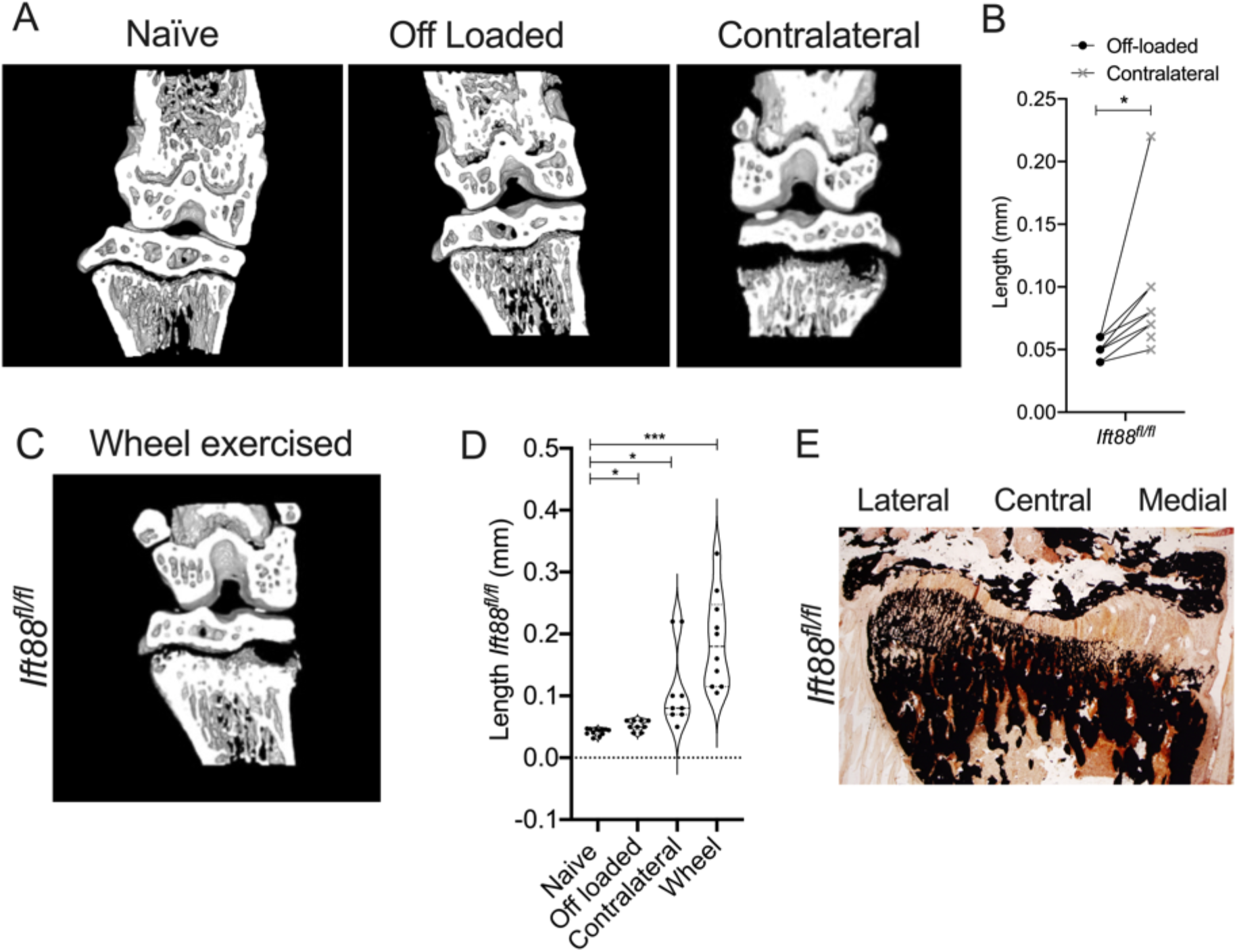
Acute increases in physiological limb loading inhibit peripheral growth plate dynamics. **A,** MicroCT partial 3D construction of off-loaded and contralateral joints from Ift88^fl/fl^ control mice. **B,** GP lengths of paired off-loaded (right) and contralateral (left) joints in Ift88^fl/fl^ control mice. **C,** MicroCT partial 3D construction of joints from Ift88^fl/fl^ control mice following two weeks wheel exercise. **D,** Quantitation of GP length of naïve, off-loaded, contralateral and wheel exercised Ift88^fl/fl^ control mice. Points in violin plots represent mean growth plate length per animal. Analysed by one-way ANOVA ANOVA, *p<0.05, ****p<0.0001, n=9-12. **E,** Von Kossa staining of Ift88^fl/fl^ following 2 weeks wheel exercise.

### Limb immobilisation rescues growth plate ossification in IFT88cKO mice

In contrast to the initial hypothesis that primary cilia may be a positive regulator of the GP response to mechanical stress, we next tested if ciliary IFT88 could be regulating GP closure, in a mechanically-dependent manner, by protecting GP dynamics from disruptive mechanical force. *AggrecanCreER^T2^;Ift88^fl/fl^* mice also underwent double neurectomy surgery to off-load the joint. In *AggrecanCreER^T2^;Ift88^fl/fl^* mice, GP dynamics were rescued in by off-loading (Figure. 4A) and the difference in GP length between genotype was abolished by off-loading (Figure 4C). The removal of ciliary IFT88 in conditions of increased mechanical loading (contralateral and wheel) did not influence the effects of increased loading on GP closure.

**Figure 4.**
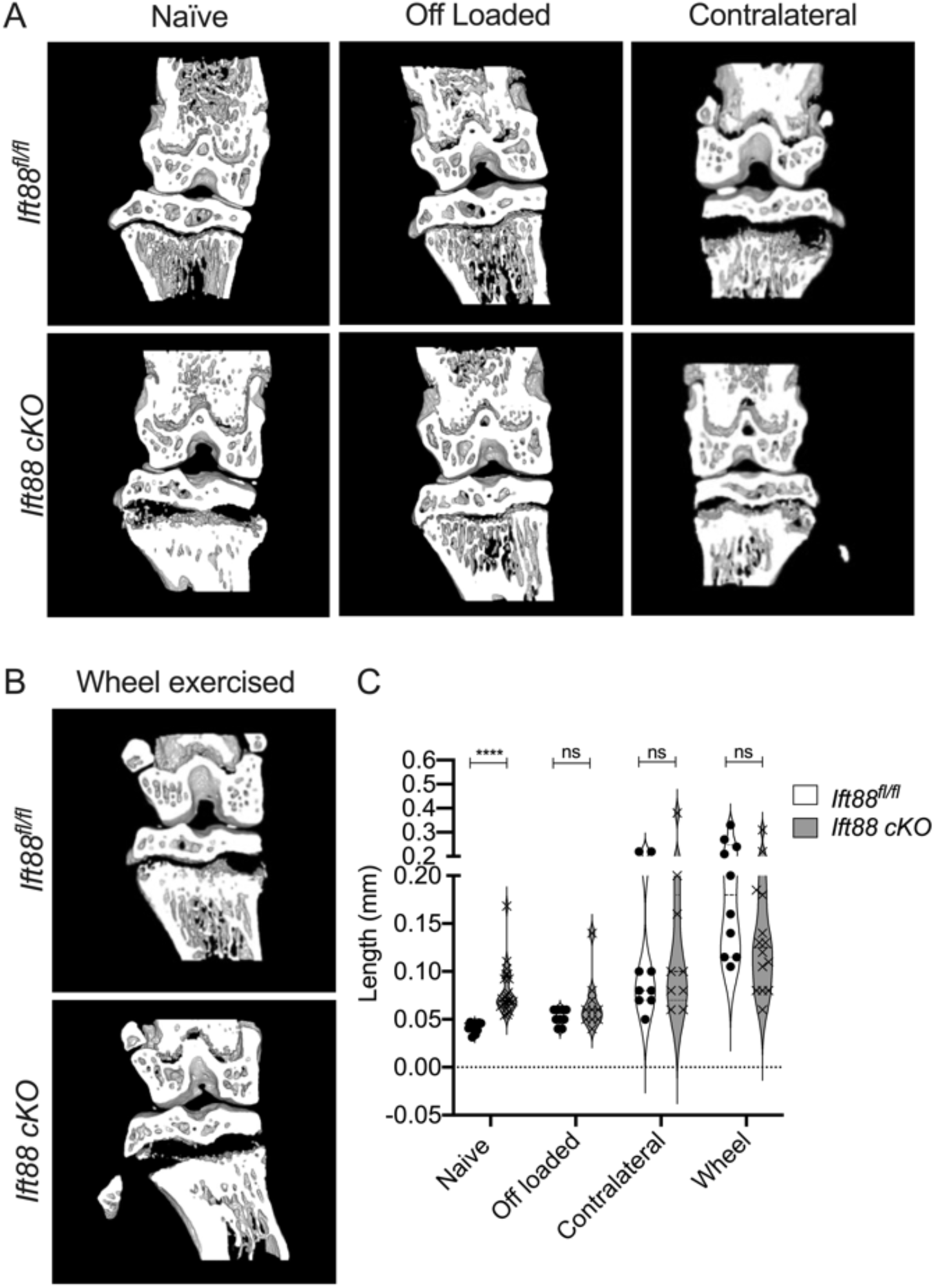
Limb immobilisation rescues the effect of IFT88 deletion on GP ossification. **A,** MicroCT partial 3D construction of AggrecanCreER^T2^;Ift88^fl/fl^ mice of naïve, off-loaded and contralateral joints. **B** MicroCT partial 3D construction of control and AggrecanCreER^T2^;Ift88^fl/fl^ wheel exercised joints. **C,** GP length of control and AggrecanCreER^T2^;Ift88^fl/fl^ mice in naïve, off-loaded, contralateral and wheel exercised mice. Points in violin plots represent mean GP length per animal. Statistical comparisons represent unpaired t-tests, corrected for multiplicity, FDR 1%, ****p<0.0001, n=9-23.

Collectively, these data indicate that IFT88 is a mechanical force-dependent regulator of the adolescent GP, apparently acting to dampen its responsivity and/or protect against otherwise disruptive physiological forces within the GP, to ensure coordinated ossification across the width of the limb. Next, we explored the cellular and molecular basis to these findings.

### Deletion of ciliary IFT88 does not impair chondrocyte differentiation, but inhibits cartilage resorption, ‘trapping’ differentiated hypertrophic chondrocytes in expanded regions of the peripheral growth plate

In order to understand the cellular and molecular mechanism underpinning the phenotype of *AggrecanCreER^T2^;Ift88^fl/fl^* mice, we assessed the cellular and matrix composition of the peripheral regions of failed ossification by histology. Safranin O staining, in naive *Ift88^fl/fl^* (control) animals revealed highly organised columns of chondrocytes in a small resting/proliferative population and larger hypertrophic population within the proteoglycan-rich GP. In contrast in *AggrecanCreER^T2^;Ift88^fl/fl^* animals, the disrupted peripheral regions directly beneath articular cartilage surfaces (Figure 5A, dashed lines), were expanded regions of proteoglycan-rich cartilage predominantly populated with swollen and disorganised, hypertrophic chondrocytes (Figure. 5A). Collagen X staining of the matrix surrounding large, hypertrophic cells indicated normal differentiation of this population (right-hand panels of Figure 5, IgG control shown in Supplementary Figure 3B). Off-loaded limbs exhibited similar GP morphology in both *Ift88^fl/fl^* control and *AggrecanCreER^T2^;Ift88^fl/fl^*, however Collagen X expression appeared reduced in both genotypes (Figure. 5B). In contrast, GP from contralateral limbs were disrupted in a manner similar to naïve *AggrecanCreER^T2^;Ift88^fl/fl^* mice, an effect only enhanced in contralateral limbs of *AggrecanCreER^T2^;Ift88^fl/fl^* where hypertrophic chondrocytes appeared even more enlarged (Supplementary Figure. 3A). Wheel exercise also resulted in enlarged peripheral regions of cartilage full of disorganised hypertrophic chondrocytes, although, as seen in immobilised limbs, Collagen X staining was weaker. This impaired ossification phenotype, observed in *AggrecanCreER^T2^;Ift88^fl/fl^* mice during adolescence, is in stark contrast to that seen with disruption of Hh signalling in embryonic and early post-natal mice, which results in accelerated hypertrophic differentiation and a reduced proliferation zone, resulting in premature ossification (29, 32, 33). Assessment of the relative populations of GP chondrocytes revealed no statistically significant changes in any population in *AggrecanCreER^T2^;Ift88^fl/fl^* mice (Supplementary Figure. 3C). However, especially on the medial side, trends towards reductions in non-hypertrophic cells, associated with increases in hypertrophic populations, were observed with deletion of IFT88 and increased limb loading (wheel exercise). This indicated, in contrast to Hh disruption, a relative expansion of the hypertrophic populations. TUNEL staining revealed very low levels, and no differences, in apoptosis, between control and *AggrecanCreER^T2^;Ift88^fl/fl^* animals (Supplementary Figure. 3D) indicating the phenotype was not associated with an inhibition of cell death at the junction between GP cartilage and bone. Thus, in *AggrecanCreER^T2^;Ift88^fl/fl^* mice, chondrocyte differentiation appeared uncoupled from GP ossification.

**Figure 5.**
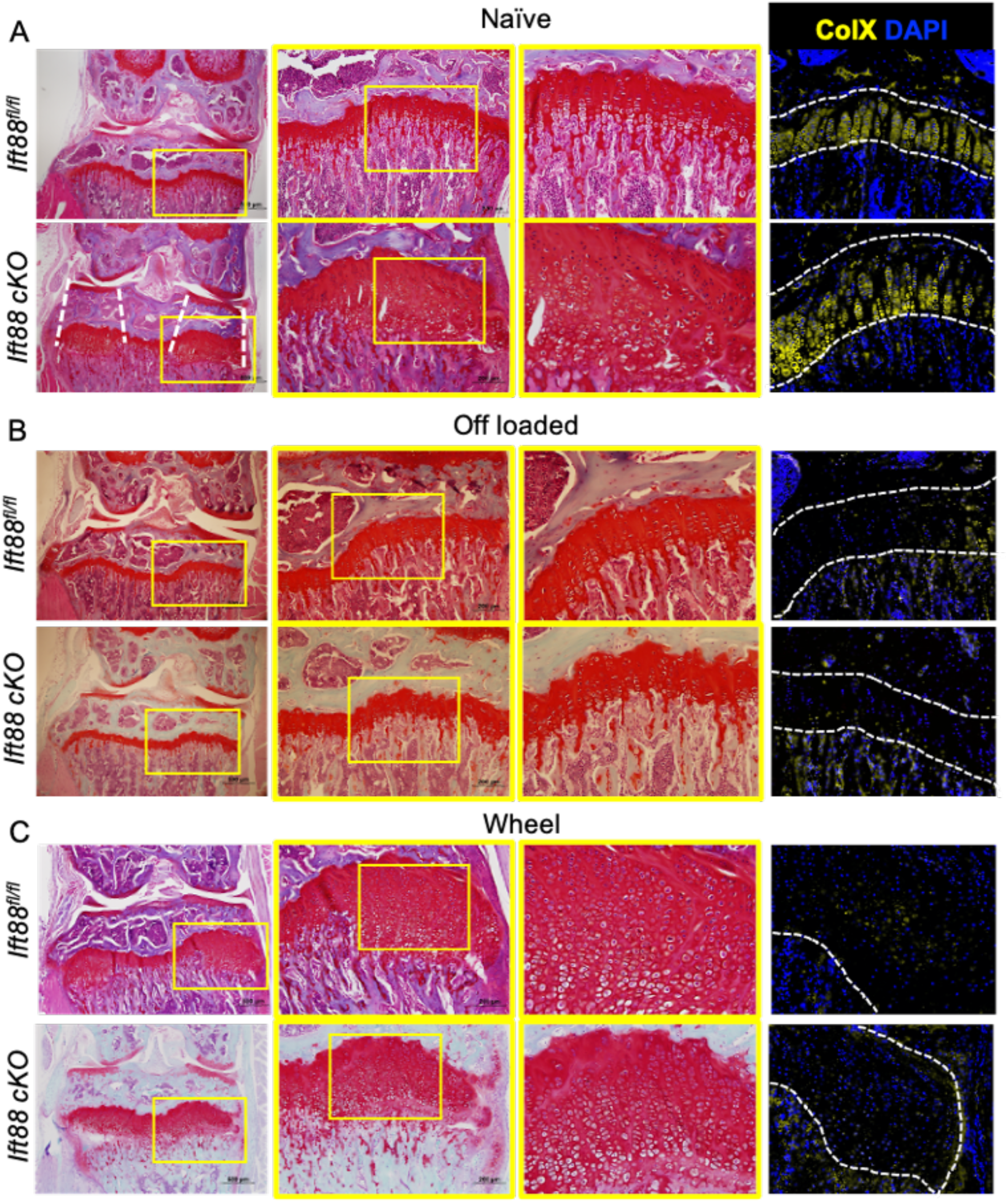
Impaired GP dynamics with IFT88 deletion and increased limb loading are associated with peripheral cartilaginous regions filled with disorganised hypertrophic chondrocytes. **A,** Safranin O stained knee joints of naïve joints. Yellow boxes on 4x (left) and 10x (middle) images are enlarged to show GP, (scale bar= 500μm). White dotted lines highlight region of GP affected is directly beneath the articular surfaces. **B,** Safranin O stained knee joints of off-loaded joints. Yellow boxes on 4x (left) and 10x (middle) images are enlarged to show GP, (scale bar= 500μm) **C,** Safranin O stained knee joints (4x) of wheel exercised joints. Yellow boxes on 4x (left) and 10x (middle) images are enlarged to show GP, (scale bar= 500μm). Representative images shown: n=6-15 in all groups. AggrecanCreER^T2^;Ift88^fl/fl^. **A, B & C,** Analysis by immunohistochemistry to assess ColX protein expression. Counterstained with nuclear DAPI. White dashed lines outline the GP, (n=5 in all groups).

To directly evaluate whether deletion of IFT88 altered GP hedgehog signalling, RNA*Scope* was performed to assess the expression of the Hh transcription factor *Gli1*, an indicator of pathway activity. *Gli1* expression was assessed on an individual cell basis and revealed that deletion of IFT88 was associated with small (13%) increases in *Gli1* expression as assessed by number of *Gli1*-positive cells (Supplementary Figure 4, ****p<0.0001, n= 4 animals in each group). This increase was most predominant in the non-hypertrophic chondrocytes (Supplementary Figure 4C). No differences in *Gli1* expression were observed when comparing peripheral regions to central regions suggesting changes to GP Hh signalling were not the primary cause of changes to GP dynamics in the peripheral regions upon deletion of IFT88.

### Deletion of ciliary IFT88 impairs osteoclastic recruitment to the peripheral growth plate

Enlarged growth plates are characteristic of protease knockout models (47, 48). It is still debated which cell types and proteases are responsible for GP resorption but we first assessed chondroclastic and osteoclastic activity at the GP/bone frontier using Tartrate-resistant acid phosphatase (TRAP) staining. In naïve, *Ift88^fl/fl^* control mice, uniform clastic activity was observed along the chondro-osseous junction (Figure 6A, top left, black arrows). In contrast, in naive *AggrecanCreER^T2^;Ift88^fl/fl^* TRAP staining was absent in peripheral regions of failed ossification (white arrows, Figure 6A), whereas the central region was largely unaffected (black arrows). In off-loaded *Ift88^fl/fl^* control joints, osteoclastic activity was reduced, in the periosteum and trabeculae, but was still present along the chondro-osseous junction across the width of the GP. In off-loaded *AggrecanCreER^T2^;Ift88^fl/fl^* mice uniform osteoclastic activity was observed across the GP thus rescuing differences between genotypes (Figure. 6A, top and bottom middle, black dotted arrows). Wheel exercise in control *Ift88^fl/fl^* mice, resulted in similar osteoclastic activity to that observed in naïve *AggrecanCreER^T2^;Ift88^fl/fl^* mice, namely a loss of TRAP staining in peripheral GP regions but not the central region (Figure. 6A, top and bottom right, black and white arrows). In exercised *AggrecanCreER^T2^;Ift88^fl/fl^,* TRAP staining was absent from the chondro-osseous junction across the width of the limb, but staining was more pronounced in the trabeculae below. Upon examination of the bone marrow at higher magnification, what appeared to be enucleated erythrocytes were visible in controls (Fig. 7B, left hand image, white arrows), at the chondro-osseous frontier. These cells appeared to invade the remnant spaces left behind by a hypertrophic cell (Figure. 6B, left hand image, black arrow). Conversely, in *AggrecanCreER^T2^;Ift88^fl/fl^* mice, there were far fewer erythrocytes and lack of bone marrow (Figure. 6B, right hand image, white arrows). Erythrocytes appeared unable to reach the growth plate to invade hypertrophic cell remnant shells (Figure. 6B, right hand image, black arrows).

**Figure 6.**
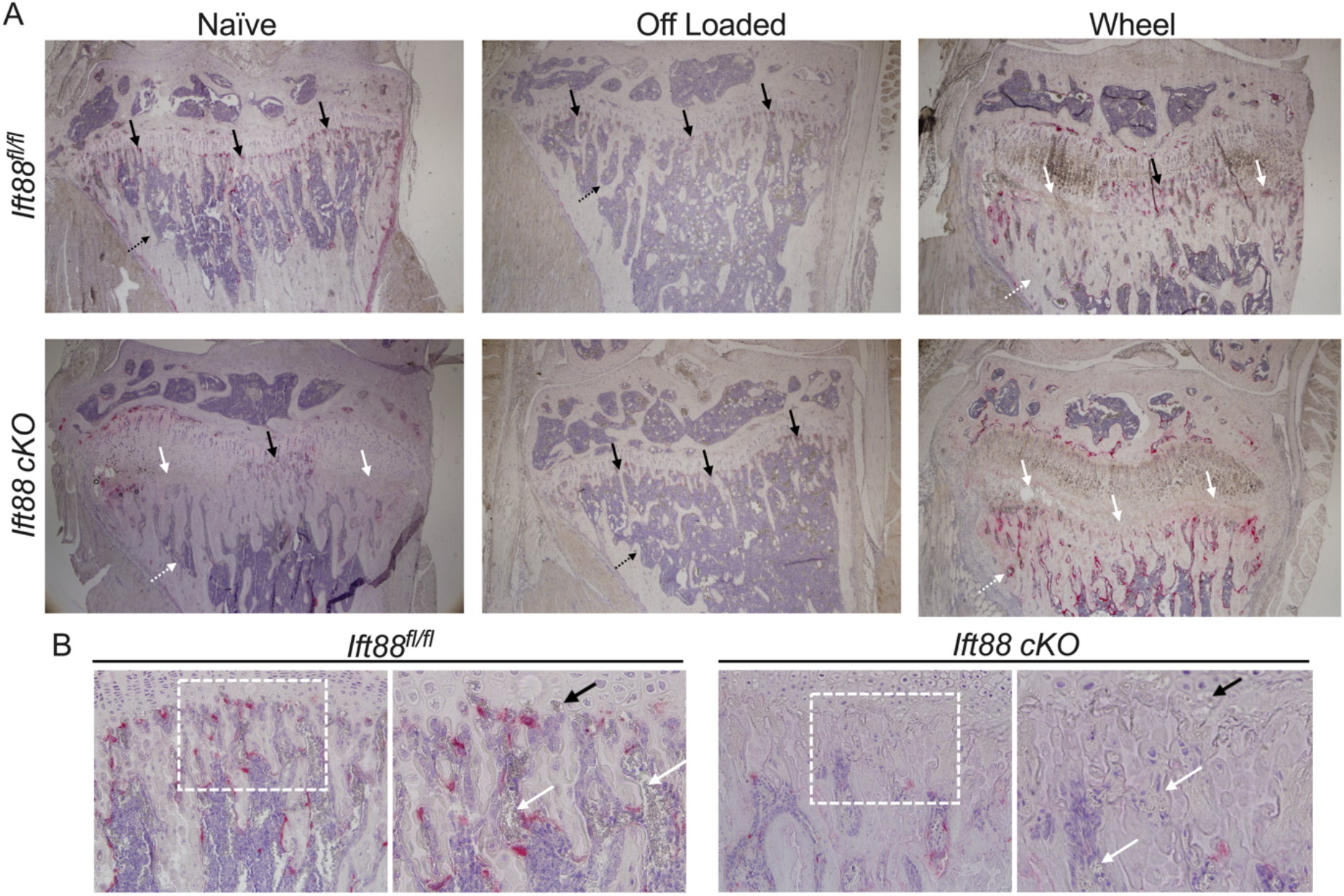
Wheel exercise and IFT88 deletion impairs osteoclast recruitment associated with failed ossification. **A,** Representative TRAP and haemotoxylin staining in naïve, off-loaded and wheel joints. Black arrows point to normal osteoclastic activity in the primary spongiosa, whereas white arrows indicate where this staining is perturbed. Black dotted arrows point to normal trabecular bone, whereas white dotted arrows point to disrupted trabecular bone formation. (n=5 in all groups). **B,** Naïve control and AggrecanCreER^T2^;Ift88^fl/fl^ mice 20x images. White boxes show area zoomed in adjacent picture. White arrows show red blood cells in the bone marrow. Black arrows point to hypertrophic chondrocyte lacunae.

**Figure. 7.**
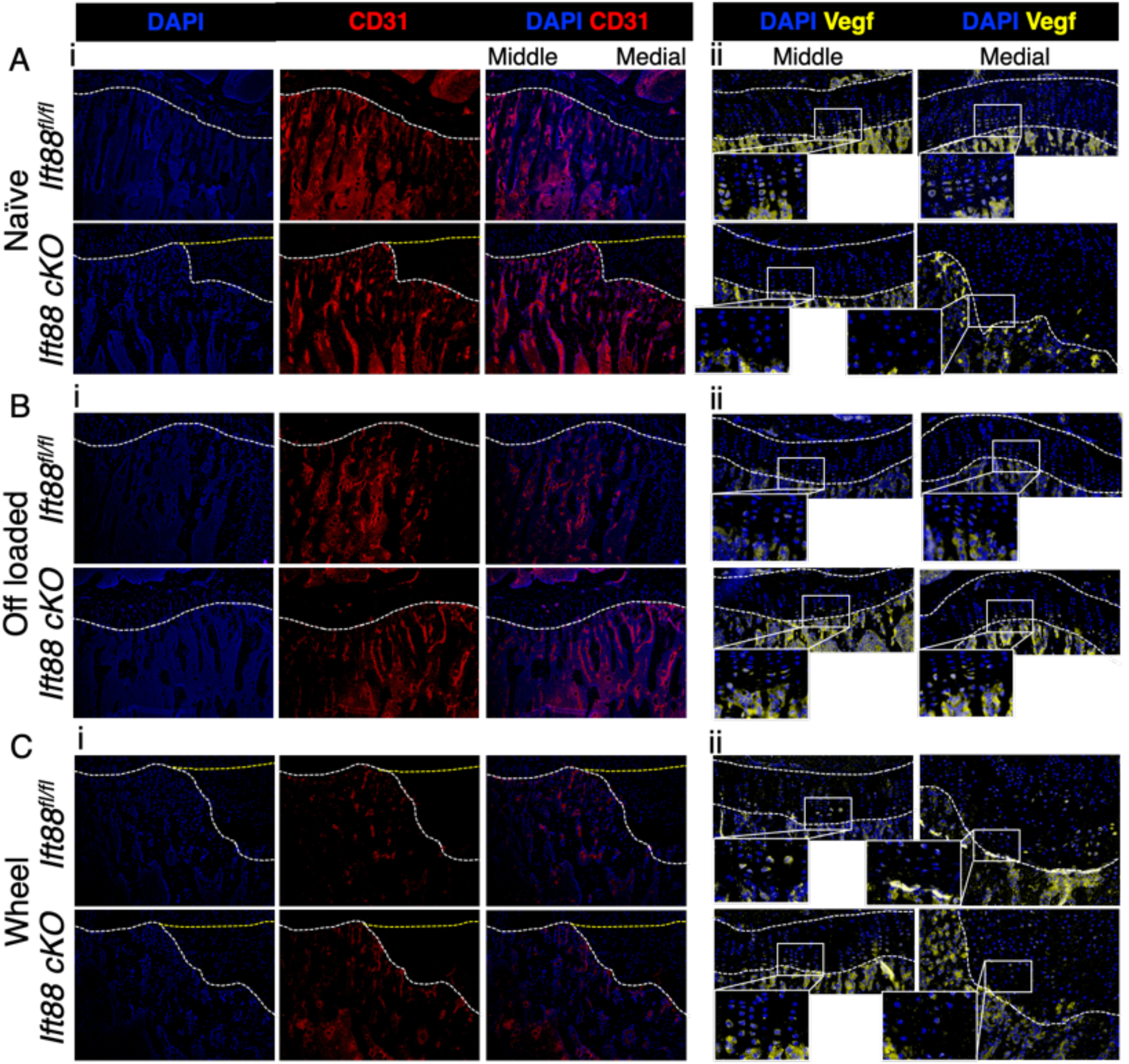
Ciliary IFT88 protects mechanosensitive expression of VEGF in hypertrophic chondrocytes. **Ai, Bi and Ci,** Representative (10x) CD31staining (red) counterstained with nuclear stain DAPI (blue) in naïve (Ai), Off loaded (Bi) and wheel exercised (Ci) Ift88^fl/fl^ control and AggrecanCreER^T2^;Ift88^fl/fl^ animals. White dashed lines demarcate the osteochondral junction between bone and GP cartilage. Yellow dashed lines indicate presumptive frontier of vascularisation if not disrupted. **Aii, Bii and Cii,** Representative (20x) VEGF staining (yellow) counterstained with DAPI (blue) in naïve (Aii), off loaded (Bii) and wheel exercised (Cii) control and AggrecanCreER^T2^;Ift88^fl/fl^ animals. White dashed lines demarcate the GP.

### Ciliary IFT88 safeguards mechanosensitive VEGF expression, enabling the coordinated vascular invasion supporting epiphyseal ossification

The carefully coordinated invasion of novel blood vessel types shapes limb development and has been shown to be critical to GP resorption during growth (49). Immunohistochemical staining of CD31, a blood vessel marker, revealed homogenous expression throughout the bone up to the osteochondral junction in *Ift88^fl/fl^* control animals (Figure 7Ai top panels). In contrast, peripheral regions of the GP cartilage where ossification failed in *AggrecanCreER^T2^;Ift88^fl/fl^*, revealed vessels were absent in these areas (Figure 7Ai bottom panels). Off-loading the joints rescued vessel invasion in *AggrecanCreER^T2^;Ift88^fl/fl^* with CD31 expression consistent across the width of the tibia in both genotypes (Figure 7Bi). However, in both *Ift88^fl/fl^* control and *AggrecanCreER^T2^;Ift88^fl/fl^* wheel exercised mice, failed regions of ossification were again associated with inhibited vascular recruitment in peripheral regions (Figure 7Ci).

To coordinate vessel invasion of the epiphyseal cartilage, vascular endothelial growth factor (VEGF) is released by hypertrophic chondrocytes. The genetic deletion of VEGF is also associated with enlarged GP (50). Histological sections were assessed for VEGF expression by IHC, revealing expression of VEGF in hypertrophic chondrocytes closest to bone in 10-week-old naïve *Ift88^fl/fl^* control mice and very strong staining in the Primary Spongiosa below (Figure 7Aii, top panels). In contrast, *AggrecanCreER^T2^;Ift88^fl/fl^* mice expressed no VEGF in regions of failed ossification, and only very low expression in the middle regions of the joint (Figure 7Aii, bottom panels). VEGF expression at the GP/bone frontier was present in off-loaded *AggrecanCreER^T2^;Ift88^fl/fl^* mice, thus immobilisation restored uniform VEGF expression across the width of the limb at the chondro-osseous junction. VEGF expression was reduced with off-loading in control mice, suggesting its expression is both IFT88 regulated and mechanosensitive. Wheel exercise in both control and *AggrecanCreER^T2^;Ift88^fl/fl^* mice, whilst not inhibitory to VEGF expression, clearly disrupted its localisation within the GP and bone beneath (Figure 7Cii). The unaffected central regions of the growth plate, expressed VEGF in a tighter localisation at the osteochondral junction.

Collectively these data indicate ciliary IFT88 regulates the mechanosensitive expression of VEGF, ensuring coordinated invasion of blood vessels, osteoclastic recruitment, GP cartilage resorption and ossification as skeletal growth draws to a close.

## Discussion

Research has repeatedly associated primary cilia with the cellular response to mechanical force, perhaps most famously in the context of propagating the kidney epithelial cell response to flow, perturbed in polycystic kidney disease (2). Given the congenital nature of the ciliopathies, the focus of cilia research has largely been on the cell and tissue development (51). Thus, our understanding of the roles of cilia in post-natal tissue has remained comparatively limited. We hypothesised that cilia would maintain influence in the juvenile and adolescent limb, where pivotal tissue adaptations follow largely pre-programmed genetic instructions, but are shaped by gradients of growth factor signalling and mechanotransduction. We have recently shown that ciliary *Ift88* is critical to the juvenile maturation and adult homeostasis of articular cartilage, controlling a program of calcification as cartilage matures (24). Here, we describe the effects of inducible and tissue-specific deletion of ciliary *Ift88* in the adolescent growth plate. We suggest the phenotype reveals how the potentially disruptive effects of mechanical forces are mitigated during this pivotal period in the limb, illustrating an example of negative regulation of the response to mechanical force by the primary cilium. We propose that, in this context, the cilium may act as a cellular and tissue ‘*mechano-dampener*’.

Analysis of a reporter line revealed a mosaic activity of the *AggrecanCreER^T2^* used to delete *Ift88*. Importantly, given the localisation of the phenotype in the IFT88cKO model (*;Ift88^fl/fl^)*, no bias was observed to Cre activity between the centre and periphery of the GP. By using IFT88*^fl/fl^* as our control, we controlled for any potential effects of tamoxifen, albeit our tamoxifen doses were below those characterised to effect bone structure (52). As we assessed tomato signal two weeks after tamoxifen administration, observed GP activity may be an underestimate of activity due to aggrecan-positive lineages transdifferentiating to the Primary Spongiosa below, although Cre activity was still apparent in resting, potentially recycling, populations at the top of the growth plate and potentially therefore active in recently described stem cell populations (25, 53). Whilst the slowing of supply of new cells to the GP may well underpin epiphyseal senescence in adolescence (26), the apparently normal progression of chondrocyte lineages within IFT88cKO GP does not suggest stem cell renewal or differentiation has been affected but rather chondrocyte GP exit is inhibited. The observation of tomato positive cells in bone is suggestive of translocation from GP to bone at this timepoint but also raises the prospect that bone resident populations have been directly affected by the Aggrecan Cre. Roles for cilia in bone progenitors (54) and osteocyte biology and mechanobiology (4) have been previously described, thus changes to the bone architecture beneath the GP may be the direct result of Cre activity on bone cell populations or indirect effects due to altered limb biomechanics or alterations in upstream ossification. Our IHC analysis was able to confirm reductions in cilia number in the GP. We cannot rule out that deletion of *Ift88* might have non-ciliary effects, including via changes to the cytoskeleton. Previous models targeting *Ift88* have documented changes to the chondrocyte cytoskeleton, implicated in regulating cellular strain (55), and more specifically in F-actin networks leading to failed hypertrophic reprogramming of chondrocytes (22). IFT proteins including IFT88 have recently been shown to interact directly with the Hippo effector YAP1 in a ciliary independent manner (56). Cartilage-specific disruption of YAP-TAZ also results in altered limb morphogenesis (57) but associated with changes in extracellular matrix.

The effects of the conditional, inducible deletion of *Ift88* we describe here, contrast with that seen in the GP of Hh and cilia models earlier in development (19, 58) (29) where Hh is a pro-proliferative signal which when lost results in premature GP closure. This immediately suggested that either the roles of Hh are altered in the adolescent GP or the most important role of cilia in the GP, at this age, is not the tuning of a Hh signal, but regulation of the cell and tissue response to another external cue. The appearance of regions of failed ossification, directly beneath the load-bearing articular cartilage plateaus, implicated an anatomical heterogeneity of tissue remodelling, potentially downstream to anisotropic tissue mechanics across the width of the limb (43, 44). This led us to hypothesise that the loss of cilia was altering GP sensitivity to mechanical force, with ramifications in peripheral regions of the tibial GP that modelling suggests experience greater stresses (43, 44). Indeed, direct analysis of Hh signalling by RNA scope did not find striking changes to the intensity or pattern of *GLI1* expression, an indicator of pathway activity, that might explain the peripheral failures of ossification. Thus, our interpretation is that the primary underlying mechanism in IFT88cKO adolescent growth plates is altered mechanoadaptation, independent of the cilium’s role in tuning Hh signalling, but through disruption of VEGF signalling.

Previous studies have investigated the effects of changes to mechanical loading during growth, for example by harnessing an extra 10% body weight to chickens (59) which resulted in narrowing of the GP and enhancements of ossification and vascularisation. In contrast, the mimicry of high impact exercise in juvenile rats monitored from 4 to 12 weeks of age also limited growth, but was associated with increased GP length (60). We were surprised to find that simply the increased compensatory loading in the contralateral limb of our double neurectomy experiments or the provision of a wheel for exercise for 2 weeks, resulted in striking inhibition of peripheral ossification of the GP in control mice, that very closely resembled that seen with conditional deletion of IFT88. We would assume, these rapid effects might recover with time, and represent an acute change for relatively sedentary caged mice, but they nevertheless demonstrate the sensitivity of GP dynamics at this timepoint. This extends the rationale for more research into the effects of acute changes to biomechanics during adolescence. In addition to modelling the effects of loading in a number of animal models, cross-sectional studies of adolescents engaged in physical activity demonstrate that sporting activity is strongly associated with epiphyseal extension and hypertrophy, and the development of CAM morphology, itself a strong risk factor for hip pain and the development of osteoarthritis (OA) in humans (61). A better understanding of the interactions between mechanical loading and the maturing skeleton will only strengthen our appreciation of the risks associated and pathological processes that underlie, common pathologies such as OA, but also other mechanically-associated chondropathies.

We propose that cilia might act to dampen or threshold the cellular response to mechanical forces in the GP that might otherwise be disruptive to its coordinated closure, predisposing the limb to poor health later in life. Cilia have been proposed to play a critical role in mechanotransduction in chondrocytes and/or the GP before, on the basis of both correlations between loading and cilia prevalence *in situ* (45) and *in vitro* evidence from chondrocyte cell lines (6, 45, 62). Cilia have recently been shown to play a critical role integrating mechanical loading and force in tendon (63). Furthermore the removal of cilia in the vascular endothelium left turbulent regions of the vasculature predisposed to the formation of atherosclerotic plaques (64), perhaps another example of ‘mechanoflammation’ recently coined in the OA field (65). We have previously investigated apparent roles for ciliary proteins in the cellular response to inflammatory cues (66, 67). As such cilia appear to act at the interface between biological and biophysical programs of bioregulation in tissue, with very likely cell type and environmental specificity. Our interpretation of the *in vivo* studies presented here is that in the adolescent epiphysis, in the absence of the influence of cilia, the differential of force, and thereby likely cellular strain in the hypertrophic region, across the width of the GP, results in heterogeneity of VEGF expression, a disrupted rather than tightly controlled expression pattern at the chondro-osseous junction (Figure 8). As demonstrated previously *in vitro (68, 69)*, limb VEGF expression is mechanosensitive *in situ,* as indicated by its scattered nature in wheel exercised mice and the reductions seen in the bone with immobilisation. *AggrecanCreER^T2^;Ift88^fl/fl^* mice exhibited a loss of VEGF expression which we propose impairs the recruitment of type H vessels (49) and osteoclasts, inhibits cartilage resorption, hypertrophic chondrocyte transdifferentiation and ultimately the coordinated ossification of the GP. These effects were all rescued upon off-loading the limb by immobilisation demonstrating the requirement of physiological loading for the IFT88 phenotype. Liu et al. (70) demonstrated that deletion of the retrograde ciliary IFT80 using a Col2a1 Cre to constitutively delete in chondrocytes, impaired chondrocyte differentiation in the context of fracture healing. The authors reported reduced angiogenesis and VEGF mRNA expression in this context. VEGF function in vascularisation has also recently been linked to cilia in pancreatic islets (71) albeit in the context of ligand internalisation and downstream signalling rather than expression. Whilst VEGF expression is mechano-regulated, the nature of the mechanical stress and identity of transducing signals driving VEGF expression that IFT88 dampens the response to, remain open questions.

**Figure 8.**
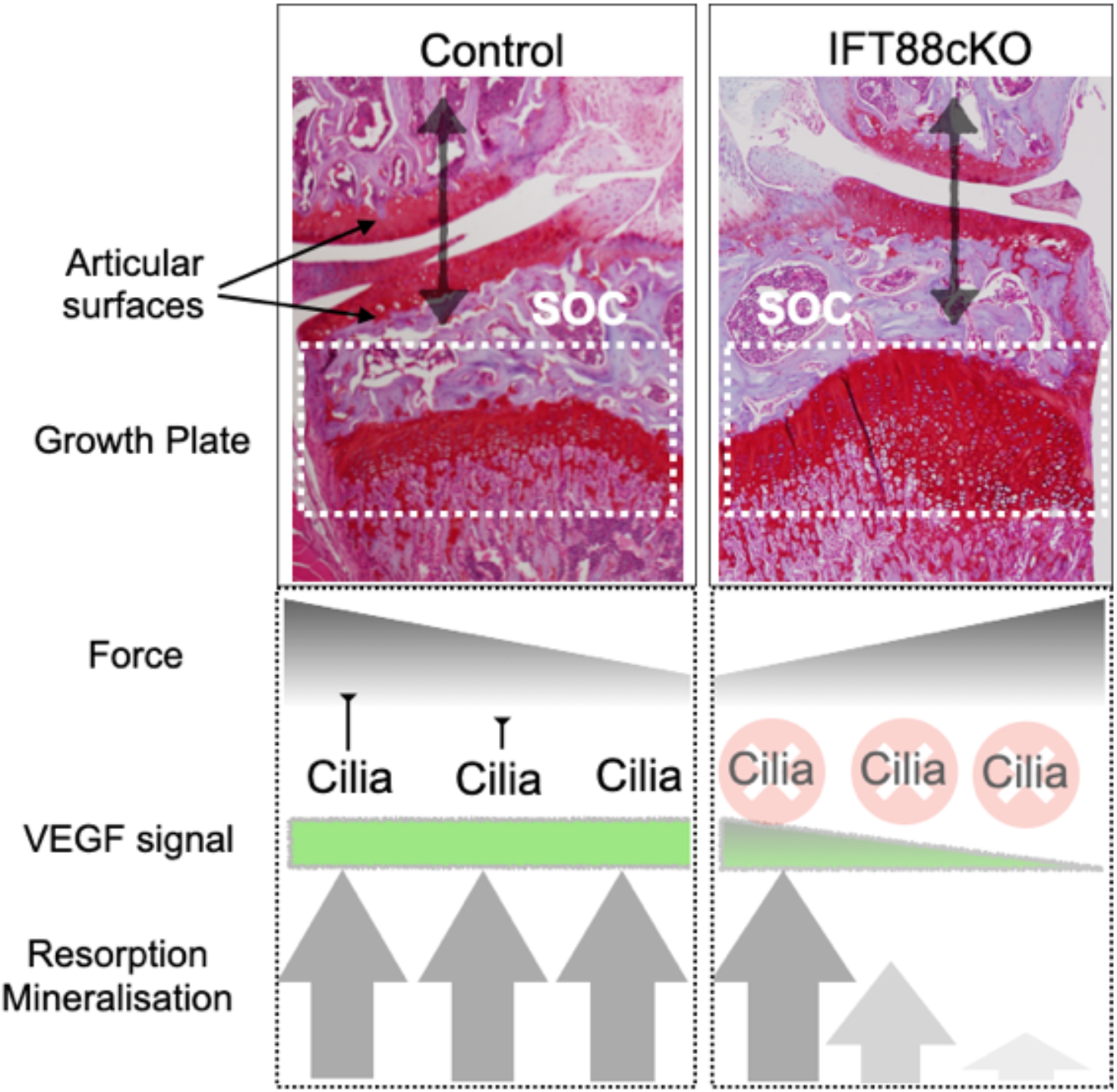
Proposed role of cilia in ensuring coordinated GP dynamics in face of disruptive force heterogeneity across the limb. In control (normal) scenario, a gradient of force exists across the width of limb, as a result of load-bearing at the articular surfaces and imperfect re-distribution of stresses by secondary ossification (SOC) centre. In this context cilia ensure an equal expression of VEGF by hypertrophic chondrocytes across the osteochondral frontier, acting to dampen responses to high loads, and potentially sensitise chondrocytes in regions experiencing lower forces. This ensures coordinated, uniform GP resorption and mineralisation. With the loss of cilia (IFT88cKO), the VEGF signal is disrupted, resulting in a failure of this coordinated advance in the peripheral regions and the ‘trapping’ of hypertrophic chondrocytes. Note: Histology images are different joints from Ift88^fl/fl^ (left) and AggrecanCreER^T2^;Ift88^fl/fl^ (right) respectively.

Both adaptability and resilience to mechanical forces are critical to tissue maturation and health. We conclude that ciliary Ift88 plays a critical role in this context in the juvenile and adolescent growth plate, its removal resulting in failed resorption and ossification of the growth plate at the end of growth. This phenomenon, observed in *AggrecanCreER^T2^;Ift88^fl/fl^* mice, is dependent on mechanical force, implying that IFT88, and potentially by extension the primary cilium, acts as a ‘m*echano-dampener*’, protecting carefully coordinated epiphyseal biology from otherwise disruptive mechanics.

## Materials and Methods

### Animals

All mice were housed in the biomedical services unit (BSU) at the Kennedy Institute, within the University of Oxford. Mice were housed 4–7 per standard, individually-ventilated cages and maintained under 12-h light/12-h dark conditions at an ambient temperature of 21°C. *Ift88^fl/fl^* mice were obtained from Jackson labs (Stock No. 022409) and maintained as the control line, and in parallel offspring were crossed with the *AggrecanCreER^T2^* mouse line, *AggrecanCreER^T2^;Ift88^fl/fl^* (Ift88 cKO), originally generated at the Kennedy Institute of Rheumatology (72). The TdTomato reporter mouse line *B6.Cg-Gt(ROSA)26Sor^tm14(CAG-TdTomato)Hze^/J* was originally from Jackson Laboratories (Stock No. 007914). For all experiments, apart from double neurectomy (off loaded) and wheel exercised (male only), both genders were used and no effect of gender was observed in the data.

### Antibodies

The following primary antibodies were used for IHC in tandem with Invitrogen AlexaFluor secondaries: Acetylated-a-tubulin (6-11B-1, MilliporeSigma, Burlington, MA, USA), Arl13b (ProteinTech, Rosemont, IL, USA, 17711-1-AP). Anti-type X collagen (polyclonal Abcam, Cambridge, MA, USA, ab58632), Anti-CD31 (Goat IgG R&D systems, AF3628) Anti-Vegf (monoclonal Abcam, Cambridge, MA, USA, ab232858).

### Tamoxifen treatment

Tamoxifen (Sigma-Aldrich, catalog no. T5648) was dissolved in 90% sunflower oil and 10% ethanol at a concentration of 20mg/ml by sonication. Tamoxifen was administered via intraperitoneal injection at ages according to experimental requirement, on three consecutive days at 50-100mg/kg (dependent on animal weight).

### Double neurectomy

One or two days prior to surgery, mice were transferred to cages containing soft bedding. Briefly, animals were prepared for surgery, anaesthesia and analgesia as previously described (73), and the right hind limb was shaved from the knee up to the hip and in the groin. Fur on the back just above the right limb is also shaved to expose the area from the spine to the flank on the right-hand side. Using a 3mm size 15 ophthalmic scalpel (MSP, Puerto Rico), a longitudinal incision is made from the right knee joint up towards and inwards towards the groin. Fine-toothed forceps were used to separate the overlying skin to reveal the muscle, femoral artery and the femoral nerve running in very close proximity. Using curved forceps, the femoral nerve is separated from its soft tissue attachments underneath. The femoral nerve is carefully transected using micro-dissecting scissors and a 0.5cm section is removed. The wound was closed using the above suturing method with additional sutures added as required. The mouse is turned on to its front and the right hindlimb is stretched out. Using a 3mm size 15 ophthalmic scalpel (MSP, Puerto Rico), an incision of approximately 2cm is made from the spine outwards. Using curved forceps, the overlying skin is separated to reveal the muscle and the sciatic nerve. The curved forceps are inserted under the sciatic nerve to separate it from the surrounding tissue. A 2-4mm region of the sciatic nerve is removed. Following this, the wound was sutured using the described method above with additional sutures added as required. Mice were transferred to a recovery chamber as described above and recovered within 10 minutes of anaesthetic withdrawal. Mice were subsequently housed in soft bedding without environment-enhancing balconies or tubes to prevent aggravation of exposed skin. Animals were disturbed as little as possible. Sudocrem was applied to the skin of the foot if aggravated. For **wheel exercise** mouse experiments, animals had access to a wheel for two weeks following tamoxifen at 8 weeks of age.

### MicroCT BV/TV

Knee joints were imaged using a MicroCT scanner (SkyScan1172 X-ray microtomograph, Antwerp, Belgium) within 70% ethanol (10μm/pixel, 3 minutes of acquisition time) (74). Using the CTan (Brucker Belgium) programme, saved image sequences were opened in the software to conduct 3D parameter analysis. Regions of interest including the epiphysis and the bone directly underneath the epiphyseal plate were defined and used to calculate the bone volume (BV), total volume (TV), ratio of BV to TV (BV/TV).

### Bridging analysis

Scans were performed with an 1172 X-Ray microtomograph (Skyscan, Kontich, Belgium). The high-resolution scans with a pixel size of 5μm were imaged. The applied X-ray voltage was 50kV, X-ray intensity 200μA with a 0.5mm aluminum filtration. The scans were taken over 180 degrees with a 0.7-degree rotation step. The images were reconstructed and binarised with a threshold of 0 to 0.16, ring artefact reduction was set at 10 using the SkyScan NRecon software package (SkyScan v 1.6.9.4, Bruker MicroCT, Kontich, Belgium). The images then were realigned vertically using DataViewer software (version 1.5.1.2 64-bit, Skysan, Belgium) to ensure similar orientation for bridging analysis. Bony bridging was analysed using a 3D quantification method as previously described (43) MicroCT scans of the tibiae were segmented using Avizo® software (V8.0, VSG, Burlington, VT, USA). The volume images were manually aligned along with the metaphyseal tibial shaft and central point of each individual bridge was selected, quantified and projected onto the tibial joint surface. From this, the areal number density of bridges (*N*, per 256 μm × 256 μm window) was then calculated, and the distribution was superimposed on the tibial surface (each bridge has a colour that represents the areal number density at the bridge location).

### Growth plate cartilage measurements

Images of histology were taken using an Olympus Osteometric microscope using a 10x lens. Quantification of cartilage width was conducted with Image J (NIH, Bethesda, MD, USA). To assess growth plate length from the lateral, medial and middle regions, maximum measurements, were taken from three consecutive sections from the middle of the joint (9 measurements per mouse). To find the length of non-hypertrophic region, the length of the hypertrophic region was taken away from growth plate length.

### Histology

Knee joints were harvested into 10% neutral buffered formalin (CellPath, Newtown, UK) for 24-48 hours. Joints were decalcified (EDTA), paraffin embedded, coronally sectioned through the entire depth of the joint. Sections (4μm), at 80μm intervals were stained with Safranin O.

### TRAP staining

70mg of Napthol AS-TR phosphate disodium salt (Sigma) dissolved in 250ul NN-dimethyl formamide (Sigma) and added to 50ml of 0.2M sodium acetate buffer at pH 5.2. 2. 115mg of sodium tartrate dihydrate (Sigma) and 70mg of fast red salt TR 1,5-napthalenedisulfonate (Sigma) was dissolved into this solution. Fixed, decalcified, unstained coronal knee sections were deparaffinised, rehydrated and placed into this solution and incubated at 37°C for 2 hours. Sections were washed briefly in deionised water and counterstained with Meyer’s Haemotoxylin (Sigma) for 1 minute and washed in deionised water before being mounted in aqueous mounting medium.

### Von Kossa staining

164μm cryosections of knee joints were defrosted in deionised water for 5 mins and incubated for 7 mins under UV light in 5% aqueous silver nitrate. Sections were rinsed thoroughly in deionised water and placed in sodium thiosulfate for 5mins, rinsed and then counterstained with Neutral red (1%) solution for 2 mins. Slides were dehydrated and mounted in Prolong Gold and visualised.

### TUNEL

*In situ* detection of apoptosis was conducted using TACS® 2 Tdt-Fluor *In Situ* apoptosis kit (Trevigen, 4812-30-K), after deparaffinising sections.

### Immunohistochemistry

Fixed, decalcified, unstained coronal knee sections were deparaffinised, rehydrated, quenched in 0.3M glycine and treated with proteinase K for 30 minutes. Samples underwent chondroitinase (0.1U) treatment for 30mins at 37°C, permeabilised by 0.2% Triton X-100 for 15mins, and blocked in 5% goat serum and 10% bovine serum albumin (BSA) in phosphate-buffered saline. Samples were incubated with primary antibody or IgG control, or no primary, overnight at 4°C. Sections were washed and incubated with Alexa-conjugated 555 secondary antibodies for 30 mins. Samples were incubated with nuclear stain DAPI (1:5000), before mounting in Prolong Gold and visualised.

### Cilia staining and confocal

Knee joints were harvested into ice cold 4% PFA and incubated in the fridge for 24 hours. Knee joints were subsequently transferred to ice cold 10% sucrose for 24 hours. This was repeated with 20% and 30% ice cold sucrose. Knee joints were then embedded into Super Cryo Embedding Medium (C-EM001, Section-lab Co. Ltd) and stored at before −80 °C. 16μm sections were collected using a pre cooled cryotome at −16 °C with Cryofilm type 3C (16UF) 2.5 cm C-FUF304. Sections were stored at −80 °C. Slides were hydrated for 5 mins in 1x phosphate-buffered saline (PBS), fixed for 10 mins with 4% formaldehyde 0.2% Triton X-100 in PBS. Sections were incubated in blocking buffer (10% bovine serum albumin, 5% goat serum in PBS) for 10 mins, followed by a 45-min incubation at RTP with primary antibody diluted in blocking buffer (1:1000 ac-a-tubulin, 1:500 Arl13b). After three 5min washes in PBS, sections were incubated with alexa-conjugated 555 secondary antibodies for 30 mins diluted in blocking buffer (1:500). After three 5mins washes in PBS, nuclei were stained using 1:5000 DAPI diluted in PBS for 5 mins, washed once in PBS and mounted in prolong gold. Imaging and analysis images were acquired using an Olympus FluoView FV1000 Confocal Microscope (Olympus, Tokyo, Japan) with an oil immersion x63 objective to produce confocal serial sections for maximum-intensity z-stack (4.6 to 5.2μm thick) reconstruction of GP sections with laser voltage, offset and gain held constant. 6 images of the growth plate per joint across the width of the tibia were taken and reconstructed. Cilia positive and cilia negative cells were blind counted by 2 individuals and their analysis averaged for each joint.

### RNAscope®

Knee joints were harvested into ice cold 4% PFA and incubated in the fridge for 24 hours. Knee joints were subsequently transferred to ice cold 10% sucrose for 24 hours. This was repeated with 20% and 30% ice cold sucrose. Knee joints were then embedded into Super Cryo Embedding Medium (C-EM001, Section-lab Co. Ltd) and stored at before −80 °C. 8μm sections were collected using a pre cooled cryotome at −16 °C with Cryofilm type 3C (16UF) 2.5 cm C-FUF304. Sections were stored at −80 °C. Slides were washed with PBS for 5mins and then baked at 60°C for 30mins. Slides were fixed using ice cold 4% PFA for 15 mins at 4 °C. Increasing concentrations of ethanol made in milli-Q water was applied, 50%, 70%, 100%, and 100% fresh ethanol, 5mins for each gradient. The sample was air dried for 5mins and incubated with hydrogen peroxide (PN 322381) for 10mins. Slides were submerged twice in milli-Q water and then transferred into pre-warmed Target Retrieval Buffer (1X) (322000) in the steamer for 10mins at 75°C. Slides were washed briefly in milli-Q water before being submerged briefly in 100% ethanol and air dried for 5mins. Protease III (PN 322381) was used to cover the sample and incubated in a HybEZ™ Oven at 40°C for 30mins. Slides were submerged briefly in milli-Q water. RNAscope® Multiplex Fluorescent Reagent Kit v2 Assay reagents (323100) was subsequently followed. We used RNAscope® Probe-Mm-Gli1-C2 (311001-C2) to assess Gli1 expression in GP cartilage and Opal™ 690 Reagent Pack FP1497001KT for visualisation. Lateral, middle and medial regions of gp were images using a 60x lens, 520nm/px, 377.6 μm × 619.77 μm, using a Zeiss 980 confocal microscope. Following normalisation with positive (RNAscope® 3-plex Positive Control Probe-Mm, PPIB gene, 320881) and negative (RNAscope® 3-plex Negative Control Probe-Mm, bacterial dapB gene, 320871) control probes, the number of Gli1 positive and negative nuclei were counted and averaged across the three regions per mouse.

## Acknowledgements

The authors wish to acknowledge Tal Arnon for provision of the ROSA26 reporter line and all members of the BSU staff at the Kennedy Institute, but particularly Albertino Bonifacio.

## Supplementary Data

**Supplementary Figure 1.**
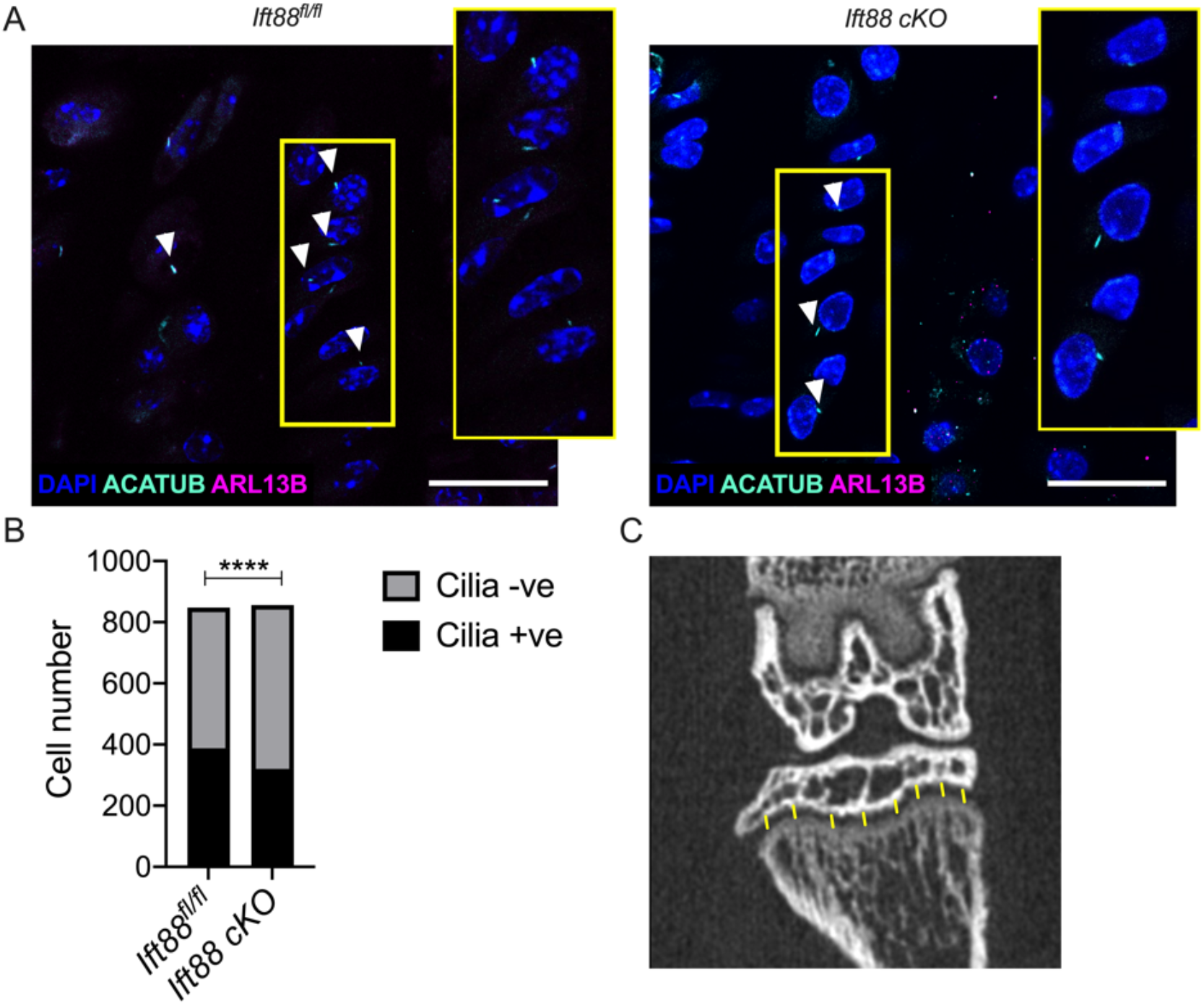
**A,** IHC staining for primary cilia in GP tissue sections from control and AggrecanCreER^T2^;Ift88^fl/fl^ animals. Scale bar 20μM. White arrows indicate clearly identifiable primary cilia positive for both Acetylated-α-tubulin (green) and ARL13B (magenta). DAPI staining (blue) indicates nuclei. **B** Cilia positive and cilia negative counts taken from 6 regions of growth plate across tibia from n=4 mice. Fisher’s exact test shown ****p<0.0001. **C** 8 points of GP length measurements (yellow lines) across representative single uCT section.

**Supplementary Figure 2.**
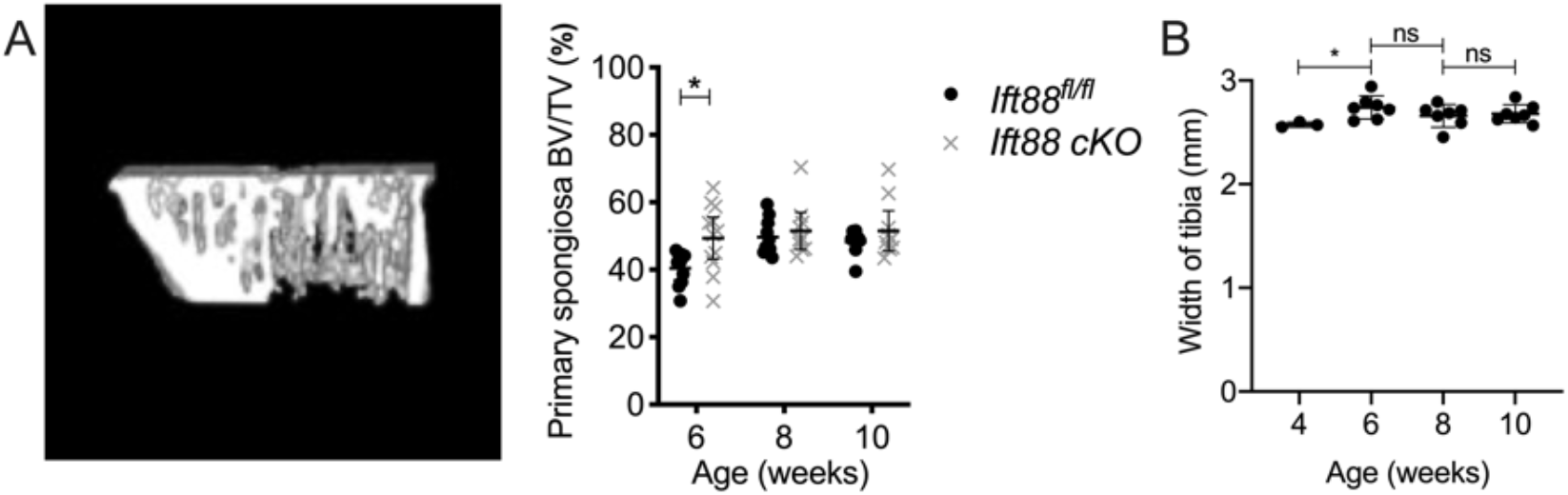
**A,** Partial 3D reconstruction of uCT scan to show the primary spongiosa region of bone directly below GP analysed and associated BV/TV (%) quantitation. **B** Tibia width measurements (from uCT). Pairwise unpaired-t-tests, corrected for multiplicity shown. *p=<0.05.

**Supplementary Figure 3.**
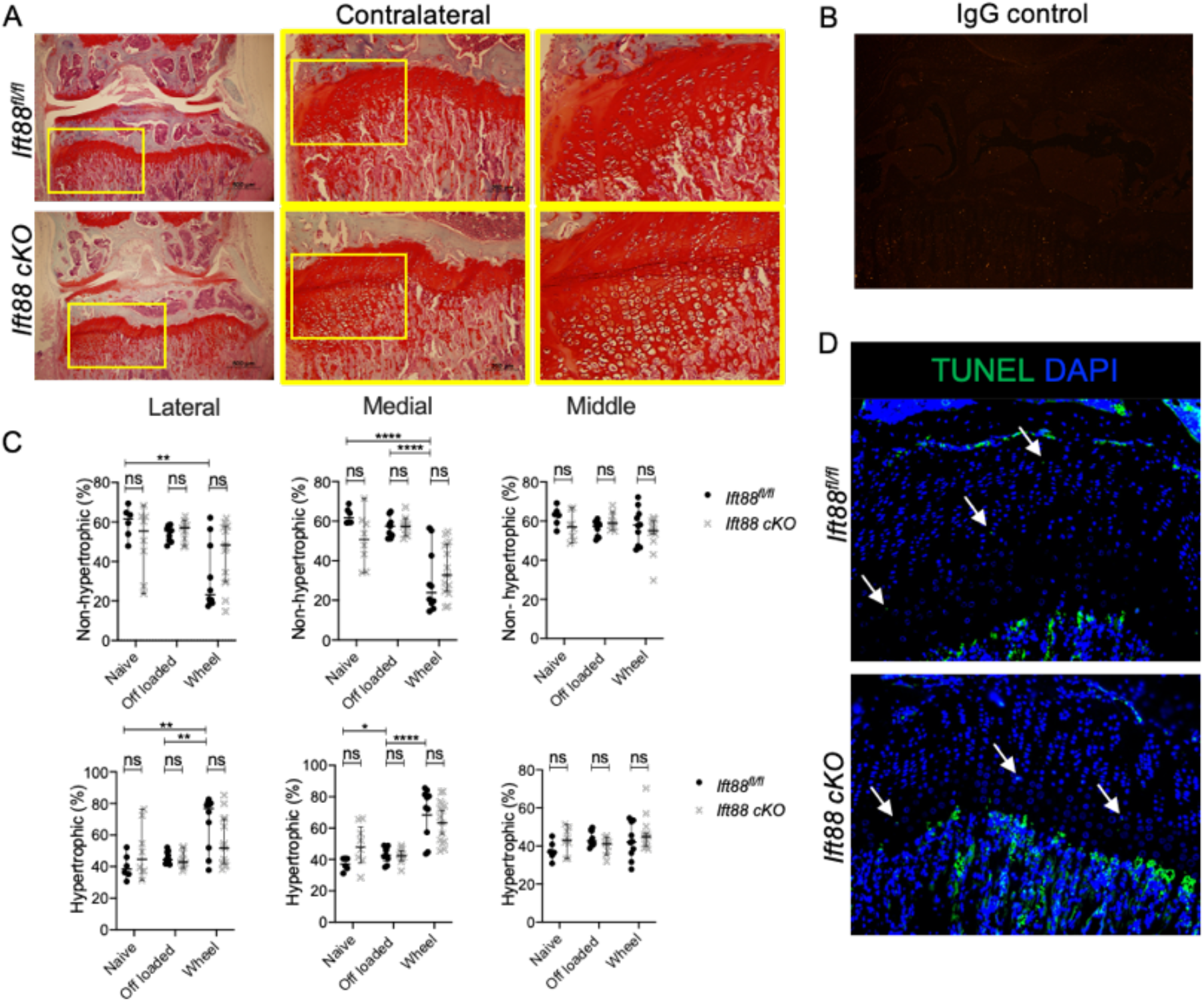
**A,** Safranin O stained knee joints of contralateral joints from control and AggrecanCreER^T2^;Ift88^fl/fl^ animals. Yellow boxes on 4x (left) and 10x (middle) images are enlarged to show GP, (scale bar= 500μm).**B,** Rabbit IgG control for conditions matched to Collagen X staining. **C** Analysis of relative (%) hypertrophic and non-hypertrophic GP chondrocyte populations. Two-way ANOVA with multiple comparison tests shown. **D** TUNEL staining (green) in control and AggrecanCreER^T2^;Ift88^fl/fl^ animals. White arrows highlight TUNEL positive cells in GP.

**Supplementary Figure 4.**
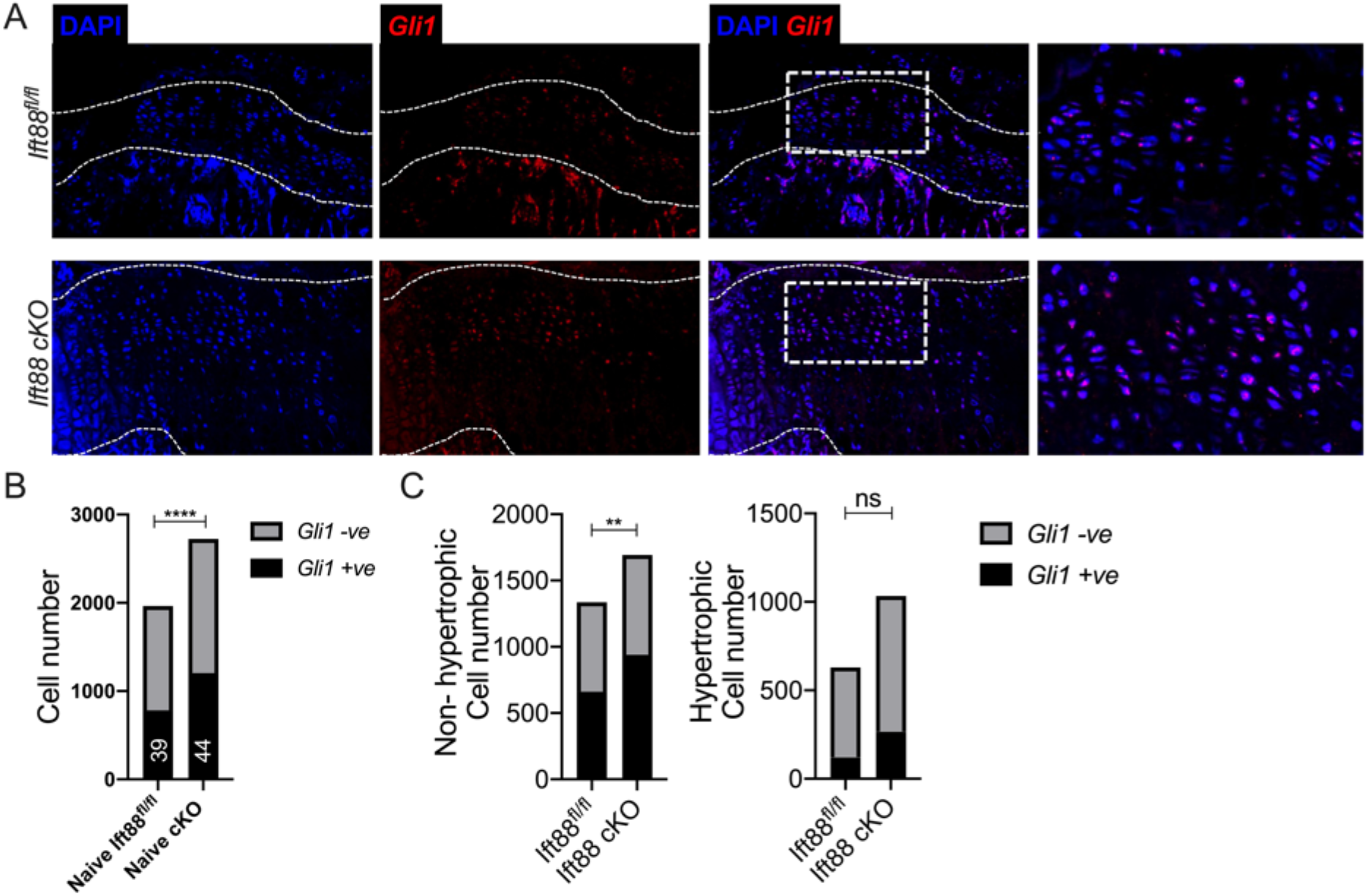
**A,** Representative RNAScope of Gli1 expression in GP on the medial side of control and AggrecanCreER^T2^;Ift88^fl/fl^ animals, counterstained with DAPI (blue) (n=4 in each group). White dashed lines demarcate GP. White dashed box shows enlarged regions in adjacent image. **B,** Contingency data of Gli1 positive nuclei (Analysed by Fisher’s exact test, ****p<0.0001, % Gli1 positive shown in white) in naïve control and AggrecanCreER^T2^;Ift88^fl/fl^ mice (n= 4 minimum in all groups). **C,** Contingency data of Gli1 positive nuclei in non-hypertrophic and hypertrophic regions of the GP to assess Gli1 expression by cell positivity (Analysed by Fisher’s exact test, **p<0.01) in naïve and AggrecanCreER^T2^;Ift88^fl/fl^ mice (n= 4 in all groups).

